# Pathogen genetics identifies avirulence/virulence loci associated with barley chromosome 6H resistance in the *Pyrenophora teres* f. *teres* – barley interaction

**DOI:** 10.1101/2023.02.10.527674

**Authors:** Jinling Li, Nathan A. Wyatt, Ryan M. Skiba, Gayan K. Kariyawasam, Jonathan K. Richards, Karl Effertz, Sajid Rehman, Robert S. Brueggeman, Timothy L. Friesen

## Abstract

Barley net form net blotch (NFNB) is a foliar disease caused by *Pyrenophora teres* f. *teres*. Barley line CIho5791, which harbors the chromosome 6H broad spectrum resistance gene *Rpt5*, displays dominant resistance to *P. teres* f. *teres*. To genetically characterize *P. teres* f. *teres* avirulence/virulence on the barley line CIho5791, we generated a *P. teres* f. *teres* mapping population using a cross between the Moroccan CIho5791-virulent isolate MorSM40-3, and the avirulent reference isolate 0-1. Genetic maps were generated for all 12 chromosomes (Ch) and quantitative trait locus (QTL) mapping identified two significant QTL associated with *P. teres* f. *teres* avirulence/virulence on CIho5791. The most significant QTL mapped to *P. teres* f. *teres* Ch1 where the virulent allele was contributed by MorSM40-3. A second QTL mapped to Ch8, however, this virulent allele was contributed by 0-1. The Ch1 and Ch8 loci accounted for 27 and 15% of the disease variation, respectively and the avirulent allele at the Ch1 locus was shown to be epistatic over the virulent allele at the Ch8 locus. Additionally, we used 177 sequenced *P. teres* f. *teres* isolates in a genome wide association study that identified the same Ch1 and Ch8 loci as the two most significant associations. Within the identified genomic regions, we identified several genes that encoded small secreted proteins, one or more of which may be responsible for overcoming the CIho5791 resistance. Results presented here demonstrate the complexity of avirulence/virulence in the *P. teres* f. *teres* - barley interaction.

## Introduction

The barley foliar disease net form net blotch (NFNB), caused by the fungal pathogen *Pyrenophora teres* f. *teres*, is a destructive disease of barley worldwide. A typical infection starts as a dot-like lesion and then develops transverse and longitudinal striations, and subsequently forms a net-like pattern of necrosis on barley leaves for which it was named (Mathre et al. 1997). Under optimal conditions, including a susceptible cultivar planted under cool and moist environmental conditions, an outbreak of NFNB can result in significant yield losses or even complete crop failure (Mathre et al. 1997; Liu et al. 2011). Furthermore, malt and feed quality of barley are negatively affected by NFNB as fungal infection leads to reduced kernel plumpness, size, and bulk density (Grewal et al. 2008; Mathre 1997). Several disease control strategies, such as foliar-applied fungicide, cultural practices, and crop rotation have been used to combat this disease, however, deployment of resistant cultivars is considered the most cost-effective and environmentally friendly method for the control of NFNB (Liu et al. 2011).

Necrotrophic fungal pathogens, especially those of the Dothideomycete class such as *Parastagonospora nodorum* and *Pyrenophora tritici-repentis*, are known to produce necrotrophic effectors (NE) that target dominant host sensitivity/susceptibility genes to induce a compatible (susceptible) interaction (Faris and Friesen 2020; Friesen and Faris 2021). However, other important Dothideomycete pathogens such as *Zymoseptoria tritici* and *Leptosphaeria maculans* secrete avirulent proteins that trigger dominant resistance gene pathways (Brown et al. 2015; Petit-Houdenot et al. 2019). Previous work on the *P. teres* f. *teres*-barley pathosystem has indicated the possibility for NEs that would be associated with fungal virulence and disease symptom development (Ismail et al. 2014). More recently, Liu et al. (2015) showed that a *P. teres* f. *teres* proteinaceous necrotrophic effector interacted with a sensitivity gene on barley chromosome 6H to induce NFNB disease. This sensitivity mapped to a centromeric region on barley chromosome 6H. Other studies have mapped dominant resistance and dominant susceptibility to the same 6H region, indicating that this 6H region is highly complex regarding NFNB. The 6H region potentially harbors genes for both gene-for-gene and inverse gene-for-gene interactions, involving effector recognition by the host resistant (R) gene and necrotrophic effectors targeting host susceptibility genes, respectively (Liu et al. 2015).

Numerous studies have shown that virulence profiles vary on differential barley lines using both local and global *P. teres* f. *teres* populations (Liu et al. 2012; Jalli and Robinson 2000; Jalli 2004, Fowler et al. 2017; Gupta and Loughman 2001; Akhavan et al. 2016; Wu et al. 2003). The diversity of *P. teres* f. *teres* virulence is likely a result of the variation in effector gene repertoire (Clare et al. 2020). Several bi-parental *P. teres* f. *teres* populations have been developed for the identification of genes associated with specific avirulence/virulence (Weiland et al. 1999; Lai et al. 2007; Beattie et al. 2007; Liu et al. 2011; Liu et al. 2015; Shjerve et al. 2014; Koladia et al. 2017b). Weiland et al. (1999) first used a *P. teres* f. *teres* population derived from the California isolate 15A and the Canadian isolate 0-1 to map a locus associated with a 15A derived avirulence on Harbin barley. Beattie et al. (2007) mapped an avirulent locus (*Avr_Heartland_*) on the barley cultivar Heartland by using a mapping population derived from a cross of two Canadian isolates differing for virulence on Heartland barley. Shjerve et al. (2014) generated a bi-parental pathogen population using the California *P. teres* f. *teres* isolates 15A and 6A. Isolate 15A showed virulence on Kombar barley but avirulence on Rika, whereas isolate 6A showed the reciprocal reaction, displaying avirulence on Kombar but virulence on Rika. Using the 15A × 6A mapping population, two independent quantitative trait loci (QTL), named *VK1* and *VK2*, were associated with 15A-derived virulence on Kombar barley, and two independent QTL, designated *VR1* and *VR2*, were associated with 6A-derived virulence on Rika barley. Each of these virulences were shown to target the same 6H region reported multiple times previously (Shjerve et al. 2014). More recently, genetic analysis of the *P. teres* f. *teres* population derived from the Danish isolate BB25 and the US isolate FGOH04Ptt-21 identified nine QTL associated with avirulence/virulence on eight different barley lines (Koladia et al. 2017b). Among these nine QTL, three had major effects that contributed over 45% of the disease reaction type variation (R^2^ ≥ 45%), while the remaining six QTL were relatively minor (R^2^ < 20%) (Koladia et al. 2017b). Using the Australian *P. teres* f. *teres* population NB029/HRS09122, Martin et al. (2020) identified a major virulent QTL on chromosome 3 using the barley cultivar Skiff, explaining 24% of the phenotypic variation (LOD = 6.6). A major QTL was also identified on chromosome 3 on Beecher barley using the NB029/NB085 population, explaining 36% of the phenotypic variation (LOD = 12.0). So far, a total of 15 unique genetic loci in *P. teres* f. *teres* have been reported, however, the genes underlying QTL for *P. teres* f. *teres* avirulence/virulence remain largely unknown (Clare et al. 2020).

Genome-wide association studies (GWAS) provide a powerful tool for detecting genetic variants associated with phenotypic variation in a population (Sánchez-Vallet et al. 2018). Due to the increased affordability of genome sequencing for fungal pathogens, GWAS has been effectively applied to characterize avirulence/virulence in several plant pathogens (Gao et al. 2016; Talas and McDonald 2015; Zhong et al. 2017; Hartmann et al. 2017; Praz et al. 2016; Bourras et al. 2019; Richards et al. 2019; Martin et al. 2020; Kariyawasam et al. 2022; Richards et al. 2022). In a recent *P. teres* f. *teres* study, Martin et al. (2020) used GWAS to identify 14 different genomic regions associated with *P. teres* f. *teres* virulence across different Australian barley cultivars and four of these genomic loci were also confirmed by bi-parental QTL mapping studies. These studies highlight the power and utility of GWAS to efficiently discover genetic variants associated with complex traits in pathogen populations.

Genetic studies using diverse barley cultivars and *P. teres* f. *teres* isolates collected from around the world have mapped both dominant resistance and dominant susceptibility to *P. teres* f. *teres* on barley chromosomes 3H and 6H, although minor quantitative trait loci have also been identified throughout the genome (Friesen et al. 2006; Abu Qamar et al. 2008, Liu et al. 2015; Shjerve et al. 2014; Koladia et al. 2017a; Clare et al. 2020).

The barley line CIho5791 (hereafter referred to as CI5791), an Ethiopian breeding line, is highly resistant to *P. teres* f. *teres* isolates collected worldwide (Mode and Schaller 1958; Tekauz 1990; Steffenson and Webster 1992; Wu et al. 2003; Cromey and Parkes 2003; Koladia et al. 2017a; Celik Oguz and Karakaya 2017). NFNB resistance conferred by CI5791 mapped to barley chromosomes 3H and 6H using the CI5791 × Tifang recombinant inbred line population (Koladia et al. 2017). It has been shown that the 6H resistance harbored by CI5791 was dominant and was effective against nine geographically diverse isolates used in the Koladia et al. (2017a) study. Intriguingly, the 3H resistance conferred by CI5791 was effective only against two Japanese *P. teres* f. *teres* isolates, whereas resistance to Danish, Brazilian, and Californian isolates mapped to a similar position on chromosome 3H with resistance being conferred by Tifang (Koladia et al. 2017a). This would indicate that either allelic variation of a single resistance gene or two closely linked resistance gene(s) is/are present on 3H in CI5791 and Tifang (Koladia et al. 2017a).

Recent studies have shown that CI5791 resistance has been overcome by a small number of select Canadian (1 isolate), French (2 isolates), and Turkish (1 isolate) *P. teres* f. *teres* isolates (Arabi et al. 1992; Akhavan et al. 2016; Celik-Oguz and Karakaya 2017). In this study, we employed QTL mapping using a bi-parental *P. teres* f. *teres* population segregating for virulence on CI5791 and a GWAS using a global *P. teres* f. *teres* population to identify genomic loci in *P. teres* f. *teres* contributing to avirulence/virulence on CI5791.

## Materials and methods

### *P. teres* f. *teres* bi-parental population development

*P. teres* f. *teres* isolate 0-1 collected from Ontario, Canada (Tekauz 1990) and isolate MorSM40-3 (hereafter referred to as Mor40-3) collected from Abda, Morocco (this study) were used to generate progeny isolates using methods described in Koladia et al. (2017b). Briefly, to set up fungal crosses, conidial suspensions of Mor40-3 and 0-1 (4000 spores/mL) were prepared and 200 μL of an equally mixed conidial suspension was pipetted onto opposite ends of sterilized wheat or barley stems that were embedded in Sach’s media (1 g CaNO_3_, 0.25 g MgSO_4_.7H_2_O, trace FeCl_3_, 0.25 g K_2_HPO, 4 g CaCO_3_, 20 g Agar, ddH_2_O to 1 L). Plates were stored in the dark at 13 °C until fruiting bodies began to develop on the stems. After approximately 3 months, mature fruiting bodies had emerged, stems were then moved to the lids of the water agar plates with the media facing opposite the stems. The plates were placed in a 12 h light/dark cycle at 13 °C. The presence of ascospores was monitored under a dissecting microscope and individual ascospores were then carefully picked from the water agar and plated on V8-PDA media (150 mL V8 juice, 10 g Difco PDA, 3 g CaCO_3_, 10 g agar, ddH_2_O up to 1 L). Single conidia were obtained from these isolated ascospore colonies. Two rounds of single spore isolation were conducted to ensure genetic purity of each progeny isolate (Shjerve et al. 2014).

### Bi-parental progeny genotyping

High-molecular weight DNA was extracted from 103 Mor40-3 × 0-1 progeny isolates following the methods described in Wyatt et al. (2018). Fungal genomic DNA was then fragmented mechanically using a Sonic Dismembrator (Fisher Scientific Part No. FB-705) using a run time of 28 minutes with a 10 second “ON” and 20 second “OFF” cycle at an amplitude of 75% of maximum. Whole genome sequencing libraries were prepared using the NEBNext Ultra II Library kit (New England Biolabs, E7645L) according to the manufacturer’s protocol. NEBNext Multiplex Oligos for Illumina (New England Biolabs, E7335L) were used to uniquely index libraries. The libraries were sequenced at Novogene Corporation (USA) on an Illumina NovaSeq 6000 platform.

### Bi-parental population marker development

Sequencing reads were quality checked using FastQC (Andrews 2010). Sequencing reads were trimmed using Trimmomatic v0.36 (Bolger et al. 2014) and trimmed reads were then aligned to the reference genome sequence of 0-1 (Wyatt et al. 2018) using the program BWA-MEM (Li and Durbin 2009). Single nucleotide polymorphisms (SNPs) were called in each progeny with the HaplotypeCaller tool from GATK (Van der Auwera and O’Connor 2020). SNP markers were filtered by following the GATK best practices (FS > 60, MQ < 40, MQRankSum < −12.5, QD < 2, QUAL < 30, ReadPosRankSum < −8, SOR > 3). All heterozygote genotypes were considered as missing data as *P. teres* f. *teres* is a haploid fungus. The marker set was finalized after applying a 30% missing data threshold, removing markers with a greater than 75% allelic frequency for either parental genotype, and thinning the number of markers so as not to have more than one marker in a 1,000 bp window of the genome (Danecek et al. 2011).

### Linkage map construction

The filtered SNP markers were imported into the linkage mapping program MapDisto v2.1.7 (http://mapdisto.free.fr/Download_Soft/) (Heffelfinger et al. 2017) for construction of genetic linkage maps. Linkage groups were created by using the “find groups” command with a LOD_min_ = 9.5 and an r_-max_ = 0.3. The “check inversions” and “ripple order” commands were used to obtain the best order of markers for each linkage group, and the “drop locus” function was used to remove potentially problematic markers. Co-segregating markers were identified and removed from the genetic map, retaining a single marker at each position. Linkage groups were named according to the corresponding *P. teres* f. *teres* 0-1 isolate reference chromosomes (Wyatt et al. 2018).

### QTL analysis

The QTL analysis was performed using the software program QGene v4.4.0 (https://www.qgene.org/qgene/download.php) (Joehanes and Nelson 2008). Composite interval mapping was performed to identify significant QTL with the default selection for cofactors as described in Koladia et al. (2017b). Critical logarithm of odds (LOD) for the population was calculated by conducting a test of 1000 permutations, and the critical LOD threshold value was set based on an α = 0.05 (Shjerve et al. 2014; Koladia et al. 2017b).

### Disease phenotyping

Parental isolates 0-1 and Mor40-3 and 103 progeny isolates were phenotyped for disease reaction on barley line CI5791. The barley seedlings were grown in the greenhouse for 12-14 days and the second fully expanded leaf of each seedling was gently taped flat to a 30 × 30 cm plastic board to provide a flat leaf surface for consistent inoculations (Wyatt et al. 2018).

Fungal inoculum was prepared as described by Koladia et al. (2017b). The inoculum was diluted to a final concentration of 1500 spores/ml and one drop of Tween 20 was added per 50 ml of inoculum to reduce spore clumping (Abu Qamar et al. 2008). The secondary leaf was inoculated with conidial solution of each progeny isolate using a pressurized paint sprayer until a heavy mist covered all the leaves but prior to runoff (Lai et al. 2007; Koladia et al. 2017b). After inoculation, plants were placed in 100% relative humidity in the light at 21 °C for 24 h and then moved into a growth chamber under a 12 h photoperiod at 21 °C for six additional days. At seven days post-inoculation, disease symptoms were evaluated on a 1–10 scale as described by Tekauz (1985), with 1 being highly resistant and 10 being highly susceptible. Three replicates were performed for each progeny isolate on CI5791 barley.

### Whole genome sequencing of a *P. teres* f. *teres* natural population

A natural population of *P. teres* f. *teres* isolates was assembled from a global isolate collection consisting of 72 isolates from the United States, 55 isolates from Morocco, 17 isolates from Australia, 21 isolates from France, and 12 isolates from Azerbaijan. Isolates were sequenced at the Beijing Genomics Institute (BGI) on an Illumina HiSeq 4000 using 150 base pair paired end reads to a target depth of approximately 30× coverage.

### Natural population marker development

The quality of sequencing reads obtained was first checked by using FastQC (Andrews 2010). Sequencing reads were trimmed using Trimmomatic v0.36 (Bolger et al. 2014) and trimmed reads were then aligned to the reference genome sequence of 0-1 (Wyatt et al. 2018) using the program BWA-MEM (Li and Durbin 2009). The alignment information was converted to binary alignment (BAM) files and indexed using the program Samtools (Li et al. 2009). Single nucleotide polymorphisms (SNPs) were called in each *P. teres* f. *teres* isolate with the HaplotypeCaller tool from GATK (Van der Auwera and O’Connor 2020). SNP markers were filtered by following the GATK best practices (FS > 60, MQ < 40, MQRankSum < −12.5, QD < 2, QUAL < 30, ReadPosRankSum < −8, SOR > 3). All heterozygote genotypes were considered as missing data as *P. teres* f. *teres* is a haploid fungus. The marker set was finalized after applying a 30% missing data threshold and a one percent minor allele frequency.

### Phenotyping of the natural population

Phenotyping of *P. teres* f. *teres* isolates from the natural population was done as previously described by Koladia et al. (2017a). Briefly, spores from each isolate in the natural population were collected in sterile distilled water and diluted to a concentration of 2,000 spores/mL with a single drop of Tween20 added per 50 mL of inoculum. The secondary leaves of barley line CI5791 were mist inoculated to just prior to runoff. Inoculated plants were placed in 100% relative humidity in the light at 21 °C for 24 h and then moved into a growth chamber under a 12 h photoperiod at 21 °C. At seven days post-inoculation, disease symptoms were evaluated on a 1–10 scale (Tekauz 1985). Three replicates were performed for each *P. teres* f. *teres* isolate on CI5791 barley.

### Genome-wide association study

A genome-wide association study (GWAS) was conducted to identify marker-trait associations for CI5791 barley. Input markers for the GWAS were first filtered for a minor allele frequency of 7%. The GWAS was performed using the GAPIT v3 software (Wang et al. 2021). Bayesian information criterion (BIC) as output in GAPIT v3 using the option ‘Model.selection=TRUE’ was used to inform the proper number of principle components for the analysis and a mixed linear model was selected, incorporating both population structure and kinship. We considered associations in the full population to be significant if *P*-values were smaller than the Bonferroni threshold (Dunn et al. 1961) at α = 0.05. A less stringent false discovery rate (FDR) (Benjamini et al. 2001) was also calculated for the global populations and for the North American and Moroccan populations alone for comparison.

### Assessment of linkage disequilibrium decay in associated loci

Linkage disequilibrium (LD) decay was calculated for the two QTL regions identified as well as for the entire chromosomes 1 and 8. Vcftools (--geno-r2) was used to obtain each pairwise comparison of LD (R^2^) between markers on Ch1 and Ch8 and pairwise LD. The decay of linkage disequilibrium with physical distance was estimated using a non-linear regression model (Remington et al. 2001; Singh et al. 2021). LD decay was defined as the pairwise distance at which LD decayed to half the maximum estimated LD for the chromosome, termed LD half decay distance. LD decay rates were compared between QTL and the respective chromosome the QTL resided on.

### Statistical analyses

Homogeneity of variance between the three replicates was performed using Bartlett’s X^2^ test. Replicates that were not significantly different at the *P* = 0.05 level were combined for QTL analysis. Genotypic classes for Mor40-3 × 0-1 progeny isolates (103 isolates) were classified by using the most significant marker associated with the QTL on Ch1 and Ch8. A Fisher’s least significant difference test was performed to identify significant variation in disease reaction type between different genotypic classes at *P* < 0.05 using data from three biological replicates using base R (R Core Team 2021).

## Results

### Bi-parental progeny phenotype analysis

Recently, a high-quality reference genome sequence was generated for *P. teres* f. *teres* isolate 0-1 (Wyatt et al. 2018), and we observed that 0-1 was avirulent on the barley line CI5791 with an average disease reaction of 1.67 on a 1 to 10 disease reaction type scale (Tekauz 1985) (Fig. 1). In contrast, Mor40-3 displayed virulence on CI5791 with an average disease reaction of 5.33 on the same scale (Fig. 1). The distinctly different virulence phenotypes for Mor40-3 and 0-1 on CI5791 led us to generate a bi-parental population from a cross between these two isolates. In total, we isolated 103 progenies derived from a sexual cross between Mor40-3 and the reference isolate 0-1 for virulence evaluation on CI5791 barley. The inoculation assays were performed three times and the disease reactions for the three replicates were homogeneous on barley line CI5791 using Bartlett’s test (*P* =0.942). The progeny isolates clearly showed a diverse range of virulence with an average disease reaction score on CI5791 ranging from 1.67 to 7.00. Transgressive segregation was present in some progenies, with three of the 103 progeny isolates exhibiting higher virulence than the parental isolate Mor40-3.

**Fig. 1.**
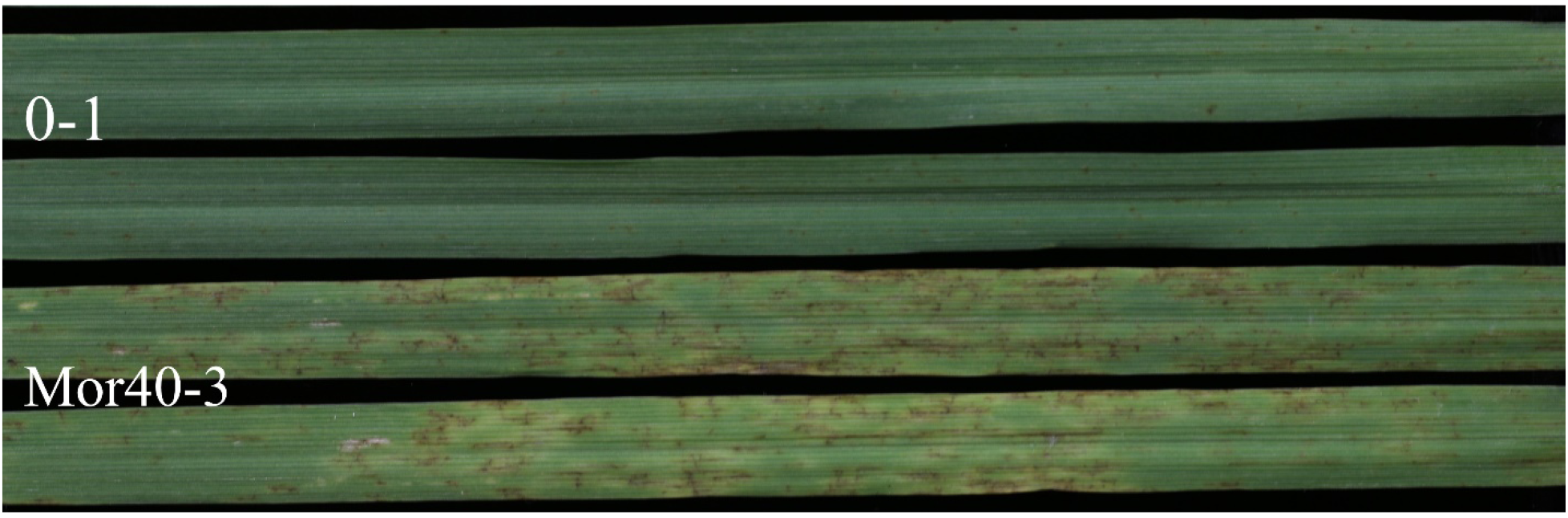
Disease symptoms caused by parental isolates Mor40-3 and 0-1 on the secondary leaves of barley line CI5791. Photo was taken 7 days post inoculation.

### Bi-parental progeny genotyping and genetic map construction

To generate a sufficiently saturated marker set for QTL analysis, we performed full genome sequencing of all 103 progeny isolates. The resulting sequencing data generated 538,873 single nucleotide polymorphism (SNP) markers with less than 30% missing data that were further filtered to 15,299 high quality SNP markers by thinning markers so that no two markers occupied the same 1,000 bp window using the parental isolate 0-1 genome as a reference (Wyatt et al. 2018). High quality markers were subsequently filtered based on allele frequency where markers possessing 25–75% of a single allelic state among the progeny were retained, and 14,226 SNP markers were exported into the linkage mapping program MapDisto v2.1.7 (Heffelfinger et al. 2017). A total of 11,995 markers were subsequently removed, as they were found to be co-segregating, were unable to be placed in any linkage group, or because they caused significant gaps, resulting in 2,231 genetically independent SNP markers being used in the final linkage maps (Table1). The resulting 12 linkage groups represented the 12 chromosomes and ranged in size from 195.32 cM to 507.85 cM with a total genetic distance of 3,842.16 cM (Table 1), spanning 92.08% of the 0-1 reference genome. (Table 1). The percentage of each chromosome covered by a single linkage group varied from 64.64% to 98.36% (Table 1). The number of non-redundant markers for each linkage group varied considerably, ranging from 106 to 296, with average marker densities ranging from a marker every 7,852 to 21,551 bp. Given that the genome size of 0-1 is approximately 46.5 Mb (Wyatt et al. 2018), the physical to genetic distance ratio in this population was approximately 12.10 kb/cM, providing a strong foundation for map-based cloning of the genes underlying the identified QTL regions.

**Table 1.**
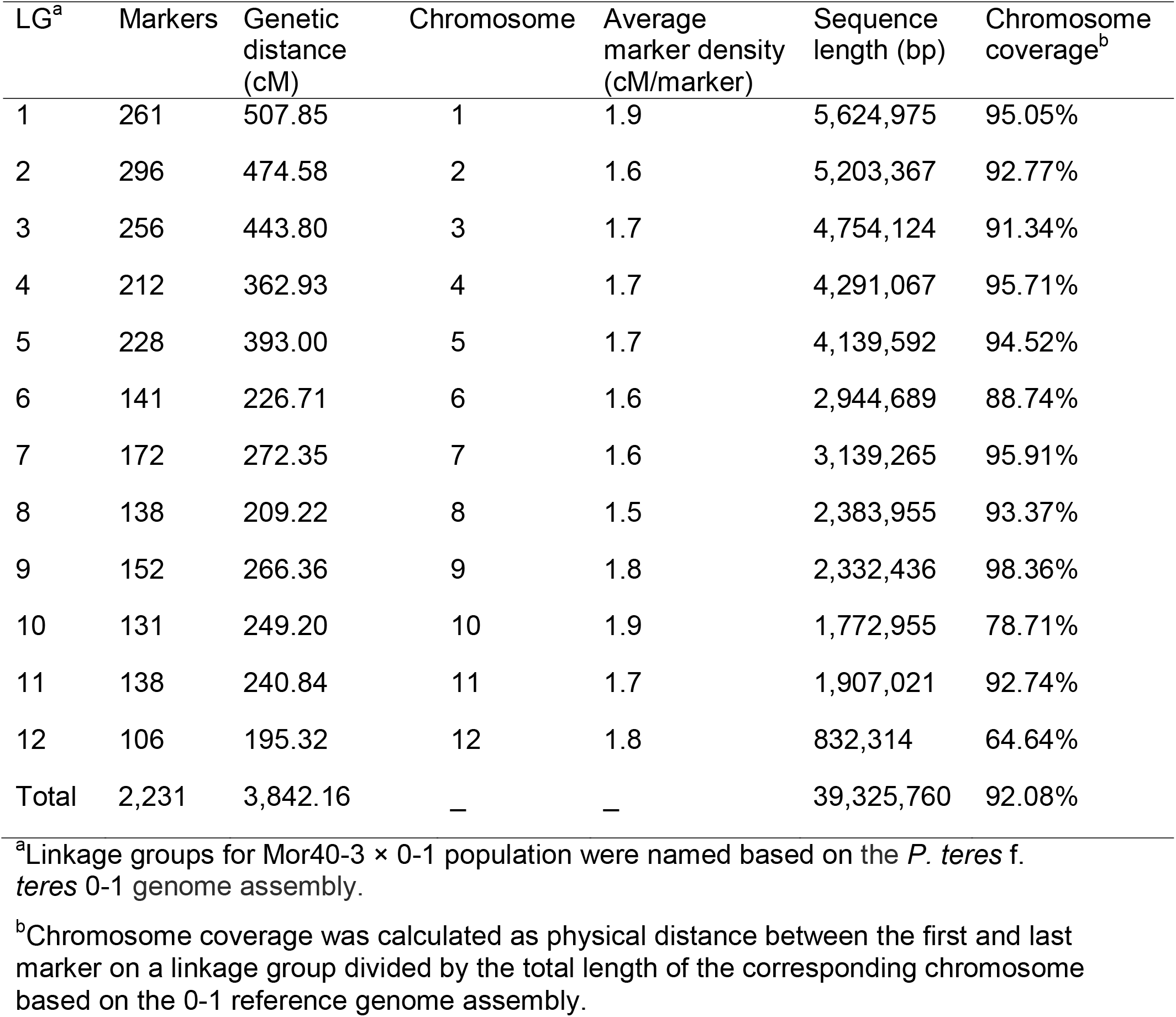
Summary statistics for the linkage groups of the Mor40-3 × 0-1 mapping population

### QTL analysis

To identify the genomic regions associated with the pathogen avirulence/virulence on CI5791, QTL analysis was performed using the genetic maps as described above and the phenotypic data collected from the 103 progeny isolates. Simple interval mapping was conducted, and a permutation test employing 1,000 permutations was used to obtain a critical LOD threshold value of 3.48 (α = 0.05) for the identification of significant QTL (Shjerve et al. 2014; Koladia et al. 2017b). Using composite interval mapping with forward co-factor selection, two major QTL were identified, one located on chromosome (Ch) 1 with a LOD value of 9.2, explaining 27% of the phenotypic variation, and the other on Ch8 with a LOD value of 7.5, explaining 15% of the phenotypic variation (Fig. 2). The virulence allele at the Ch1 QTL was contributed by parental isolate Mor40-3, while the virulence allele at the Ch8 QTL was contributed by the avirulent parent 0-1. In addition, a minor QTL (with LOD value 4.9, explaining 11% of the phenotypic variation) was identified on Chr3 with the virulent allele being contributed by Mor40-3.

**Fig. 2.**
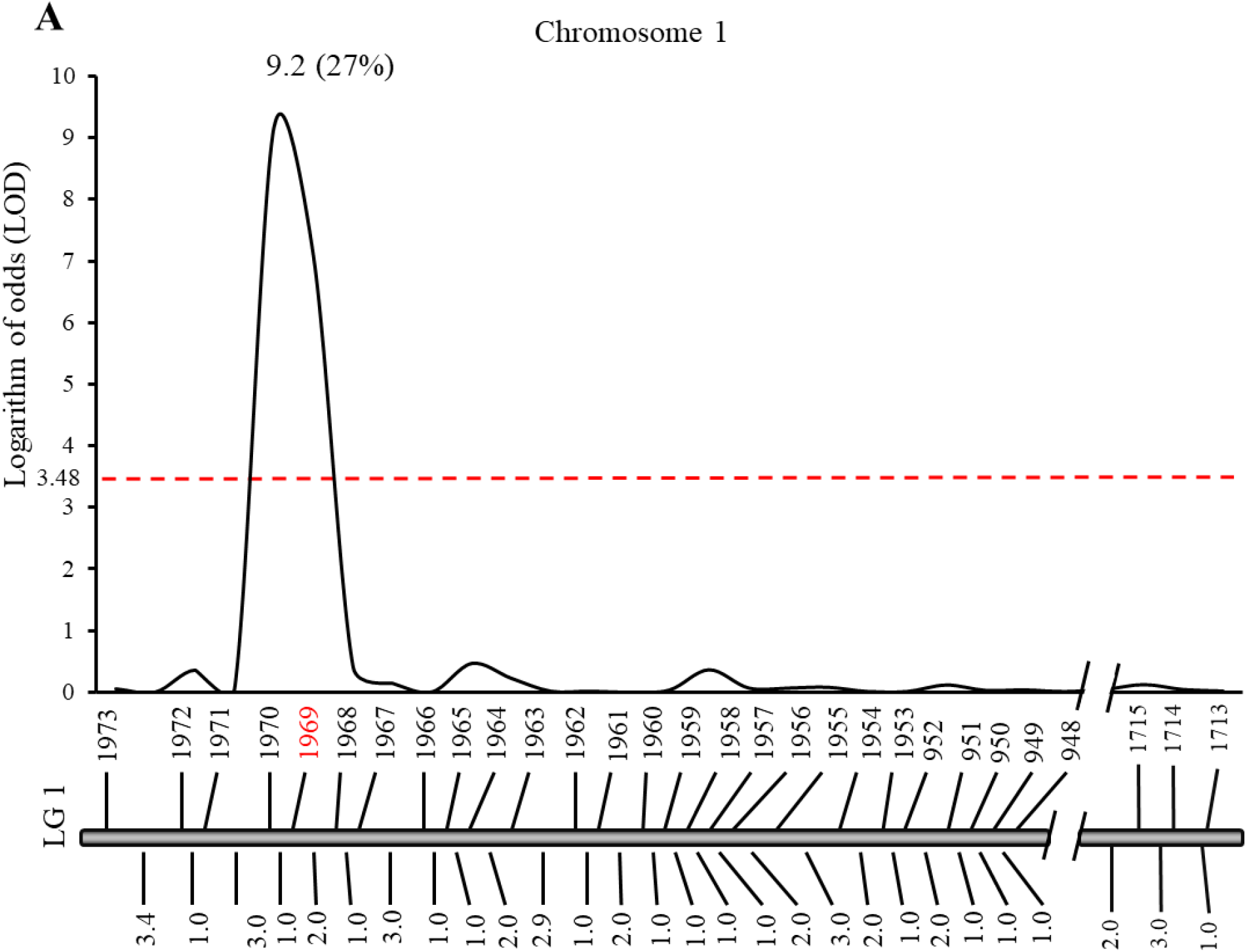
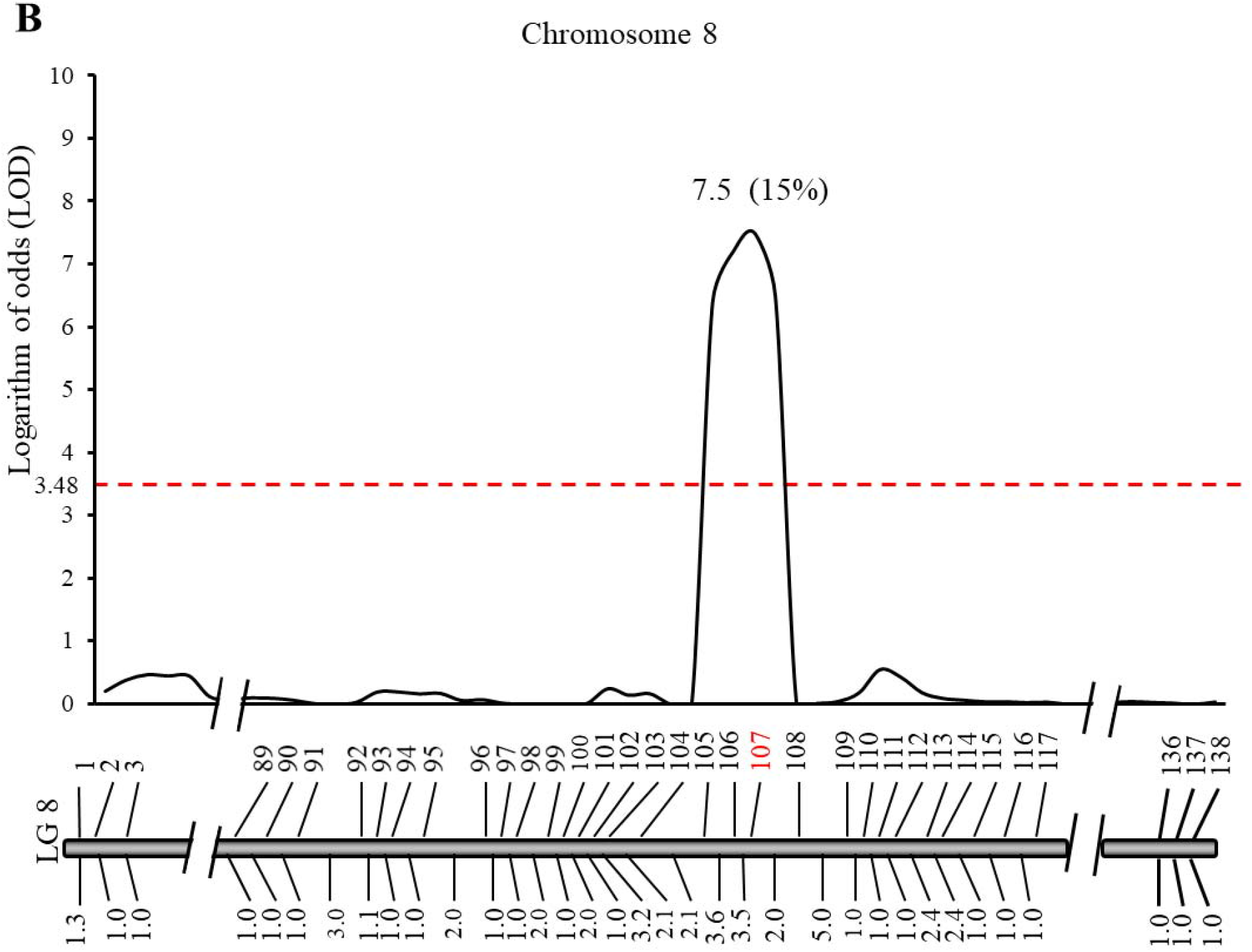
Quantitative trait loci (QTL) analysis of *P. teres* f. *teres* virulence in the Mor40-3 × 0-1 population on CI5791 barley. The x-axis shows the position of the QTL composite interval mapping regression curve on chromosomes 1 and 8. The y-axis represents LOD values with the LOD threshold of 3.48 (α = 0.05) represented by the horizontal red dashed line. The LOD and R^2^ values for each significant QTL are shown on top of their corresponding QTL peaks. The corresponding SNP markers are shown on top of the genetic maps and genetic distances on the bottom. The most significant marker for each QTL is shown in red.

To assess the effects of the Ch1 and Ch8 QTL on avirulence/virulence of isolates associated with CI5791, a Kruskal-Wallis test was used to look for significant differences between average disease scores when the progeny isolates were separated into four different genotypic groups including Ch1^Mor40-3^/Ch8^0-1^ (harboring the Mor40-3 allele at the Ch1 locus, 0-1 allele at the Ch8 locus, i.e. both virulent alleles), Ch1^Mor40-3^/Ch8^Mor40-3^ (harboring the Mor40-3 allele at both loci), Ch1^0-1^/Ch8^0-1^ (harboring the 0-1 allele at both loci) and Ch1^0-1^/Ch8^Mor40-3^ (harboring the 0-1 allele at Ch1 locus, Mor40-3 allele at the Ch8 locus, i.e. both avirulent alleles). Progeny isolates with the Ch1^Mor40-3^/Ch8^0-1^ genotype, which contained both virulence alleles, showed the highest average disease reaction, and this group was significantly higher than all other genotypic groups (Fig. 3, Table 2) indicating the importance of both loci. Progeny isolates with the Mor40-3 parental genotype Ch1^Mor40-3^/Ch8^Mor40-3^ had significantly higher average disease reactions than the other two genotypic groups (Ch1^0-1^/Ch8^0-1^, Ch1^0-1^ /Ch8^Mor40-3^) (Fig. 3, Table 2). However, the disease reactions caused by the progeny group harboring the Ch1^0-1^/Ch8^0-1^ genotype was not significantly different than the isolate group harboring the Ch1^0-1^ /Ch8^Mor40-3^ genotype (Fig. 3, Table 2), indicating that the avirulence allele (0-1 allele) at the Ch1 locus was epistatic to the virulent allele (0-1 allele) at the Ch8 locus, explaining why the 0-1 parental isolate was completely avirulent on CI5791.

**Fig. 3.**
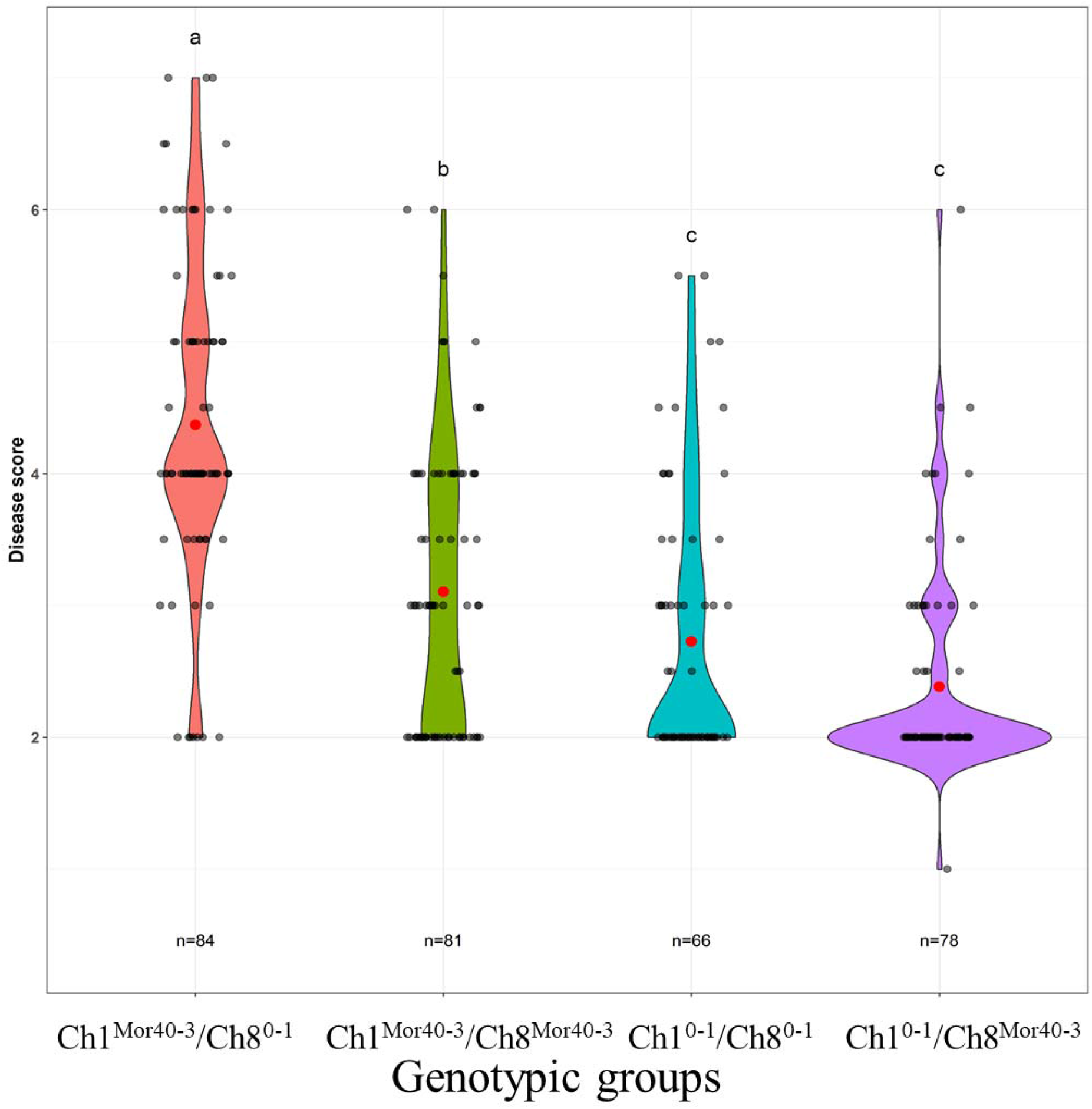
The virulence profiles of Mor40-3 × 0-1 progeny isolates from four genotypic groups. Violin plots showing the distribution of disease reactions (y-axis) of progeny isolates from the four genotypic groups (x-axis) inoculated on barley line CI5791. Red dots represent the mean disease reaction of each genotypic group, black dots represent individual disease reactions of each progeny isolate from three biological inoculations. Fisher’s least significant difference test was used to determine the significant variation in disease reaction type at the 0.05 level of probability. Disease reactions from genotypic groups followed by the same letter (above each plot) are not significantly different at the 0.05 level probability.

**Table 2.** Average disease reaction types of Mor40-3 × 0-1 progeny isolates according to genotypic class

| Genotypic class | No. of isolates | Disease range | Average disease score* |
| --- | --- | --- | --- |
| Ch1<sup>Mor40-3</sup>/Ch8<sup>0-1</sup> | 28 | 2.00-7.00 | 4.37 a |
| Ch1<sup>Mor40-3</sup>/Ch8<sup>Mor40-3</sup> | 27 | 2.00-5.33 | 3.10 b |
| Ch1<sup>0-1</sup>/Ch8<sup>0-1</sup> | 22 | 2.00-4.50 | 2.72 c |
| Ch1<sup>0-1</sup>/Ch8<sup>Mor40-3</sup> | 26 | 1.67-4.50 | 2.38 c |

\*Average disease reaction scores that are followed by different letters indicate statistical differences according to Fisher's least significant difference test ( $P = 0.05$ ).

We further mapped these two QTL to the 0-1 reference genome sequence and found that the confidence interval for the QTL on chromosome 1 spanned a 69 kb genomic region (Ch1: 1,382 - 71,035), while the QTL on chromosome 8 spanned a 148 kb genomic region (Ch8: 1,861,626 - 2,009,599). As recent studies from our lab indicated that effector proteins play an important role in the *P. teres* f. *teres-* barley pathosystem (Liu et al. 2015), we focused on genes that encoded putative secreted proteins within the identified QTL regions. A total of seven genes were found in the Ch1 QTL region, of which one was predicted to encode a secreted protein by SignalP 4.0 (Petersen et al. 2011) and this candidate gene was predicted to encode an effector by EffectorP 3.0 (Sperschneider and Dodds, 2022) (Table S1). Three of 43 genes were predicted to encode secreted proteins (Petersen et al. 2011) within the Ch8 QTL region with only one of these genes being predicted to be an effector by EffectorP 3.0 (Table S1).

### *P. teres* f. *teres* genome wide association study

To identify markers associated with virulence on CI5791 barley, a global population of 177 *P. teres* f. *teres* isolates were sequenced and markers were identified in reference to isolate 0-1 (Wyatt et al. 2018). A total of 4,249,769 SNP markers were identified and subsequently reduced to 83,828 markers after filtering for 30% missing data and a minor allele frequency of 7% to be used for the GWAS. GWAS results are summarized in Table 3. A mixed linear model was used to identify significant associations with the CI5791 phenotypic data on all chromosomes except Ch5, Ch7, Ch11 and Ch12 (Table 3, Fig. 4A) indicating a diversity of virulence factors present globally. The most significant markers identified on Ch1 were identified within the Ch1 QTL region identified above. The most significant marker identified in Ch8 was just adjacent to the QTL region identified in the bi-parental QTL study, however it was in marker disequilibrium with the markers within the QTL region, indicating it is likely identifying the same locus.

**Table 3.**
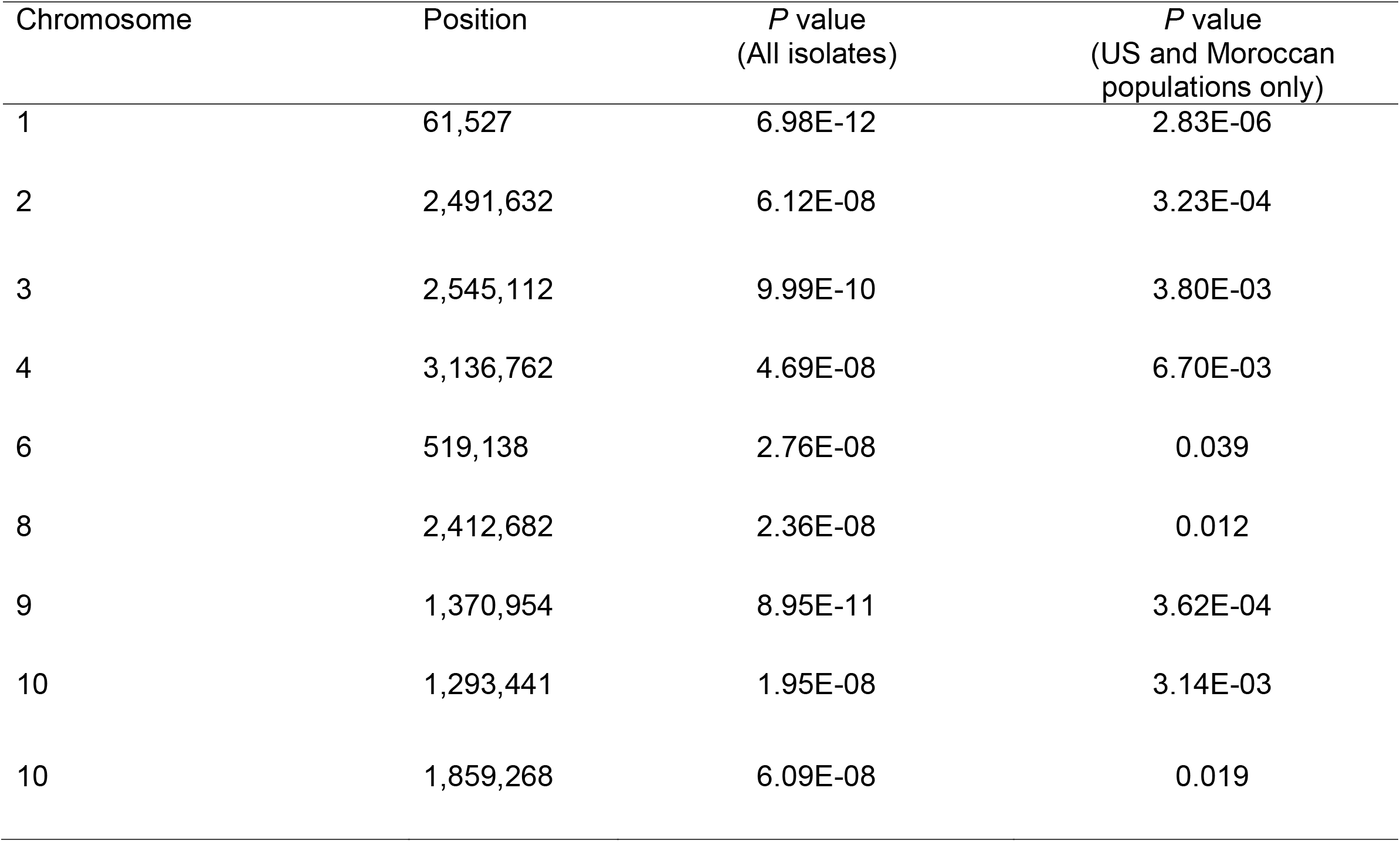
Summary of significant loci identified in the genome wide association study using the *P. teres* f. *teres* natural population.

**Fig. 4.**
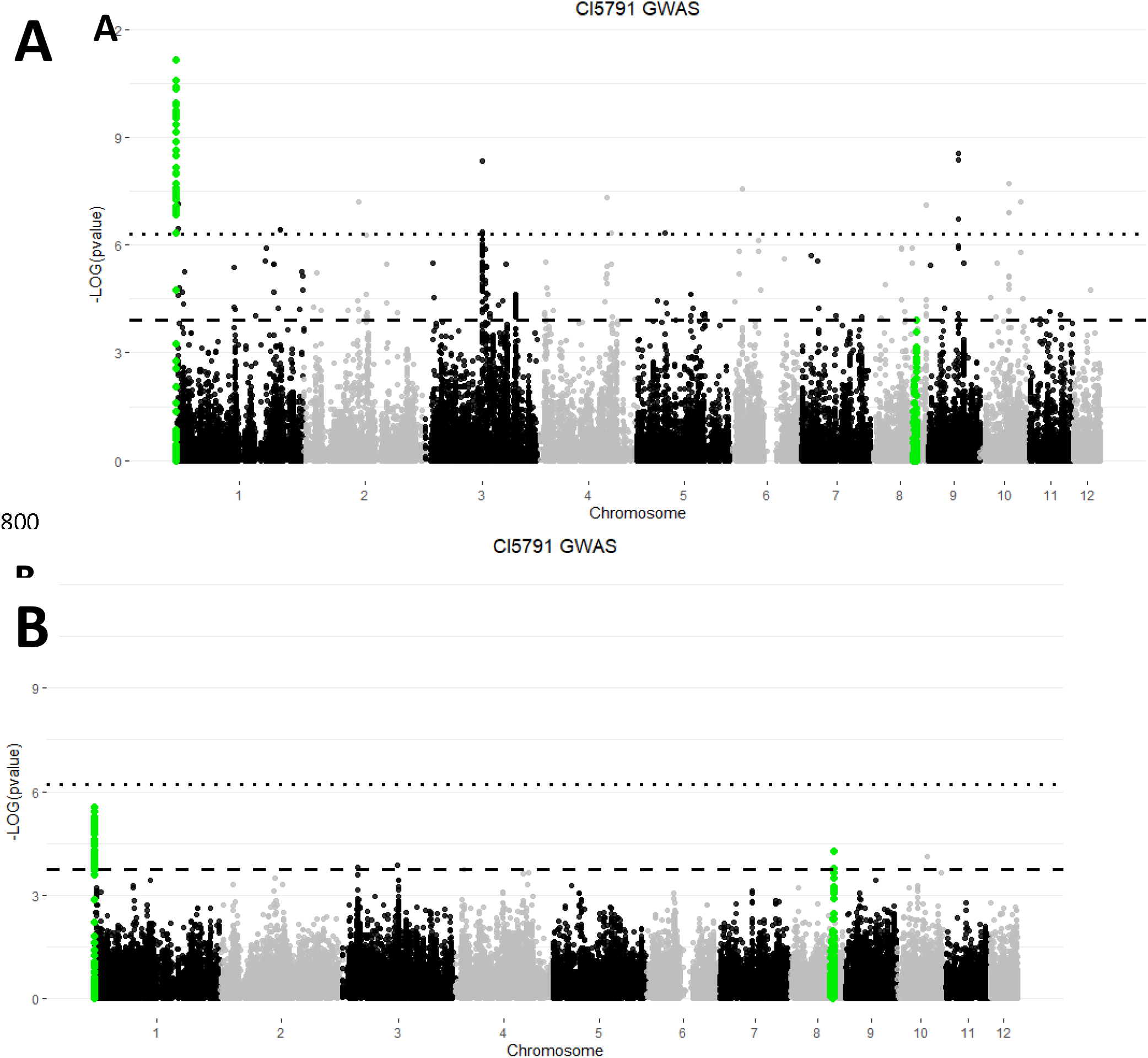
Manhattan plots representing the results of the genome wide association study (GWAS) performed using the markers generated from the *Pyrenophora teres* f. *teres* natural population and the phenotypic data collected on CI5791 barley. Plot A was generated using the entire 177 isolate global collection and Plot B was generated using data from only the isolates collected in North America (72 isolates) and Morocco (55 isolates). Each dot represents an individual marker with alternating black and grey colors representing chromosome boundaries. Markers colored green represent markers within the bounds of the quantitative trait loci (QTL) identified using the *P. teres* f. *teres* bi-parental population Mor40-3 × 0-1. The horizontal dotted lines represent the -log(p-value) Bonferroni significance thresholds of 6.3 (A) and 6.2 (B). The horizontal dashed lines represent the -log(p-value) false discovery rate (FDR) thresholds of 3.9 (A) and 3.7 (B). The most associated markers on Ch1 and 8 of the Moroccan/North American GWAS (Plot B) fell within the *P. teres* f. *teres* Ch1 and Ch8 QTL regions shown in Figure 2, however all markers fell below the level of significance in this analysis when using a Bonferroni threshold but attained significance using the FDR threshold.

To further dissect the significant associations identified for the barley line CI5791, the natural population was subset into a population that reflects the previously mentioned bi-parental mapping population Mor40-3 × 0-1. Therefore, a new GWAS dataset that was used as a mirror of the bi-parental population was generated using only isolates collected in North America (72 isolates) and Morocco (55 isolates) and evaluated for marker-trait associations using the CI5791 phenotyping data. The two most significant marker trait associations identified in this GWAS analysis were identified on Ch1 and Ch8 and were in a similar genomic region identified by the QTL analysis using the Mor40-3 × 0-1 bi-parental population (Fig. 4B), providing validation that the genes underlying these associations were important in the global population. When using the highly conservative Bonferroni correction significance threshold, neither of these loci were significant, likely due to the reduced population size. However, when using the less conservative false discovery rate (FDR) significance threshold, both the Ch1 and Ch8 loci were significantly associated with virulence on CI5791. Using the FDR, Ch3 and Ch10 also had significant associations, again indicating the potential for other virulences present in the global population. Collectively, the GWAS and QTL analyses show that the Ch1 and Ch8 loci are consistently associated with overcoming the CI5791 source of NFNB resistance.

### Assessment of linkage disequilibrium decay in associated loci

Linkage disequilibrium (LD) decay was assessed and compared between the identified QTL regions and their respective chromosomes. For Ch1, the chromosome-wide calculated distance by which LD decayed to half its maximum (R^2^=0.23) was 74,134 bp and a similar result was observed for Ch8 as LD decayed to half the maximum (R^2^=0.22) at 69,847 bp. By comparison, the QTL region of Ch1 had a LD half decay distance of 2,743 bp and the Ch8 QTL region had a LD half decay distance of 8,711 bp. Indicating more frequent recombination within the virulence-associated regions relative to the entire chromosome where they resided, a hallmark of genomic regions harboring effectors.

## Discussion

Recent work on both sides of the *P. teres* f. *teres*-barley system has shown that this host-pathogen interaction is highly complex and includes both dominant and recessive resistance on the host side and predicted effectors conferring avirulence and virulence on the pathogen side. Many of these interactions have been associated with a barley genomic region near the centromere of chromosome 6H that harbors both resistance and susceptibility genes as well as other NFNB disease associated genes (Ameen et al. 2021; Clare et al. 2020). Understanding the pathogen effectors targeting or being recognized by these 6H gene products as well as the 6H genes themselves is necessary to completely decipher the molecular interplay and evolution between *P. teres* f. *teres* and barley.

CI5791 has historically been one of the most common differential lines when characterizing global populations of *P. teres* f. *teres*, likely due to its almost universal resistance response to *P. teres* f. *teres* isolates collected across geographically diverse barley growing regions. This near universal resistance is due to the NFNB resistance locus *Rpt5* (Manninen et al. 2006). Although virulence has been identified on CI5791 in a few studies, these studies have identified only one or two virulent isolates (Arabi et al. 1992; Akhavan et al. 2016; Celik-Oguz and Karakaya 2017; Tekauz 1990) with the vast majority of global isolates being avirulent on CI5791. The rarity of virulence on CI5791 indicates that either the CI5791 resistance is not commonly used in popular cultivars planted globally or that avoiding the CI5791 resistance results in a major fitness penalty for the pathogen unless it is necessary to complete its pathogenic life cycle. Cloning of the pathogen effector gene(s) will allow us to evaluate the evolution and mode of action of this effector more closely.

Koladia et al. (2017a) showed that CI5791 harbored a dominant resistance gene at chromosome 6H that was effective against a global collection of *P. teres* f. *teres* isolates, indicating that the CI5791 6H resistance source (*Rpt5*) was a potentially durable resistance that could combat the global NFNB problem. In the adjoining paper, Richards et al (202_). showed that CI5791 gave a differential response to a natural *P. teres* f. *teres* Moroccan population and where the CI5791 resistance was effective, the resistance localized to the same Ch6H centromeric region, however, no 6H association was present when the population was inoculated with the Moroccan isolates virulent on CI5791. The dominant resistance coming from Tifang previously identified at chromosome 3H (Koladia et al. 2017a) was however, highly effective against these CI5791-virulent isolates. Richards et al. (submitted) subsequently used progeny isolates from the Mor40-3 × 0-1 population presented here that were virulent on both CI5791 and Tifang to demonstrate that progeny with virulence combinations could overcome both the 6H and 3H resistances, showing that although this virulence combination was not identified in the North American or Moroccan populations, it is possible that natural populations could overcome both dominant resistances via recombination. This recombination event emphasizes that these major resistances could be overcome if put into a cultivar and planted over large areas. These major resistance genes should be used with caution and used in conjunction with other qualitative and quantitative resistances to preserve the efficacy of these resistance genes.

The research presented here clearly showed that the Moroccan *P. teres* f. *teres* population had overcome the CI5791 broad-range dominant resistance localized to barley chromosome 6H. This discovery provided an opportunity to genetically characterize the avirulence/virulence genes interacting with the CI5791 6H gene(s). Therefore, we crossed the CI5791-virulent Moroccan isolate, Mor40-3, with the *P. teres* f. *teres* reference isolate 0-1 (Ellwood et al. 2010; Wyatt et al. 2018) that was avirulent on CI5791. We generated saturated genetic maps of all 12 *P. teres* f. *teres* chromosomes and, as a validation step, we used a GWAS approach with a global *P. teres* f. *teres* natural population. Both bi-parental and GWAS approaches identified associations with virulence at similar positions on *P. teres* f. *teres* Ch1 and Ch8.

The most significant virulence association mapped to Ch1 with the virulent allele being conferred by the virulent parent Mor40-3, however, the virulent allele at the Ch8 locus was contributed by the avirulent parent 0-1. This was interesting since 0-1 was highly avirulent (1.67 type reaction) on CI5791. When comparing genotypic classes of progeny isolates, the highest average disease reaction types were found in the progeny isolates harboring both the Ch1 and Ch8 virulent alleles (Ch1^Mor40-3^/Ch8^0-1^) followed by the genotypic class harboring only the Ch1 virulence (Ch1^Mor40-3^/Ch8^40-3^) (Fig. 3; Table 2). However, genotypic classes harboring only the Ch8 virulence allele (Ch1^0-1^/Ch8^0-1^) was not significantly different than the genotypic class harboring neither virulence allele (Ch1^0-1^/Ch8^40-3^). The significance of the Chr8 virulence only in the presence of the Ch1 virulence allele showed that the Ch1 avirulence was epistatic to the Ch8 virulence. This epistatic association between the alleles at Ch1 and Ch8 indicated that the Ch1^Mor40-3^ allele was required for the evasion of an early CI5791 recognition that resulted in an early host defense response. Given that the CI5791 6H resistance is dominant (Koladia et al. 2017) and results in early onset pinpoint lesions, our working hypothesis for the Ch8 virulence is that the underlying Ch8^0-1^ allele likely encodes for an effector with a role in facilitating colonization but is only effective after an evasion of the early defense response that requires the Mor40-3 allele at the Ch1 locus. Cloning of the genes underlying the Chr1 and Chr8 QTL is critical to the validation of this hypothesis.

An increased number of loci involved in *P. teres* f. *teres* avirulence/virulence have recently been identified through bi-parental mapping studies, which could be used for breeding resistant barley cultivars (Clare et al. 2020). The major QTL on Ch1 identified in the current study was previously identified in a different *P. teres* f. *teres* bi-parental population (Koladia et al. 2017b), which indicated the importance of these genes in the avirulence/virulence of various *P. teres* f. *teres* isolates. The major QTL on Ch8 and the minor QTL on Ch3 appear to be novel, as both were located a significant distance from any previously described locus on Ch8 or Ch3, respectively (Clare et al. 2020; Martin et al. 2020). *P. teres* isolates harboring the three QTL that were identified in this study could be used in barley breeding for resistance to *P. teres* f. *teres* in the future.

In addition to the two major Ch1 and Ch8 QTL, we also identified a Ch3 minor QTL (accounting for 11% of the disease variation), highlighting the complexity of this barley - *P. teres* f. *teres* interaction where both major and minor loci are involving in NFNB disease. Moreover, additional significant loci associated with avirulence/virulence on CI5791 were identified in the GWAS analysis on the global population on Chromosomes 2, 3, 4, 6, 9, and 10(2), indicating that in addition to the Ch1 and Ch8 associations found in the bi-parental population, several other loci that likely harbor effector genes were important across the global population but were not necessarily segregating in the bi-parental population used here.

To validate this hypothesis, a GWAS analysis was done using only the US and Moroccan populations. From this analysis, the Ch1 locus had the most significant marker-trait association and the Ch8 locus had the second most significant association, indicating that the Ch1 virulent allele was present and was the most significant contributor to CI5791 virulence in the Moroccan population and the Ch8 virulent allele is likely prevalent in the US population based on the virulent allele being present in 0-1 but not in Mor40-3. Based on the FDR threshold used in this GWAS, three other loci harboring MTAs were identified with two being on Ch3 and one on Ch10. Both Ch3 MTAs were different from those identified in the global GWAS study, however the MTA identified on Ch10 was also identified in the global GWAS (Figure 4B).

To understand why the CI5791 6H resistance is so effective and to be able to predict the durability of this gene, we must also understand how the pathogen is able to overcome this source of resistance. The genetic characterization of loci associated with virulence is the first step and this is now complete. However, the cloning, validation, and functional characterization of the underlying genes and their allelic variation is critical to a complete understanding of this source of resistance. We have already released a near-complete telomere-to-telomere reference quality assembly of 0-1 annotated with RNAseq support (Wyatt et al. 2018) and have recently generated a reference quality genome assembly of Mor40-3 (unpublished data). These resources, in conjunction with the marker-saturated Mor40-3 × 0-1 map and the GWAS data from our global population provide a great foundation for the cloning and characterization of the virulence-associated genes.

Using 0-1 as a reference, The Ch1 QTL region consisted of 69 kb and harbored seven predicted genes, one of which was predicted to be a small, secreted protein. The Ch8 QTL region consisted of 148 kb and harbored 43 genes with three of these genes predicted to encode secreted proteins. These genes will be at the top of the candidate gene list; however, more work will need to be done to annotate the Mor40-3 genome and identify the genes showing polymorphism between these two parental isolates. Given the fast LD decay in natural fungal populations (Gao et al. 2016; Richards et al. 2019), GWAS identification of marker trait associations at these loci will also be beneficial in narrowing the candidate gene regions. Given that the LD half decay of the Ch1 and Ch8 QTL regions was calculated at 2,743 bp 8,711 bp, respectively, GWAS will be a valuable tool in narrowing these genomic regions and prioritizing the effector gene candidates contributing to virulence. Identification and validation of the genes underlying these QTL/MTAs is currently underway in our lab.

## Statements and Declarations

Mention of trade names or commercial products in this publication is solely for the purpose of providing specific information and does not imply recommendation or endorsement by the U.S. Department of Agriculture. USDA is an equal opportunity provider and employer.

## Author contribution statement

TLF, RSB, JL, and NAW initiated the study and designed the experiment. JL, NAW, RMS performed experiments and data analysis. SR and KE provided fungal isolates. JL, NAW and TLF wrote the manuscript, and all authors contributed to the final version.

## Acknowledgements

This work was supported in part by the U.S. Department of Agriculture, Agricultural Research Service through project 3060-22000-051-000D.

Research was supported by the NSF/NIFA-AFRI Plant Biotic Interaction program of the National Science Foundation (award #1759030) and the USDA, National Institute of Food and Agriculture (award #2018-67014-28491), and the USAID Linkage Program.

## Conflict of interest

The authors declare that they have no conflicts of interest.

## Data Availability

The raw sequencing reads of a natural population of *Pyrenophora teres* f. *teres* are available at the National Center for Biotechnology Information (NCBI) GenBank under BioProject PRJNA923641. The raw sequencing reads of the bi-parental *Pyrenophora teres* f. *teres* population are available at NCBI GenBank under BioProject PRJNA923640.

## Supplementary material

**Table S1.** Summary of predicted effector candidate genes within quantitative trait loci (QTL) genomic regions

| Gene | SignalP<br>4.0<br>(Y/N)<sup>a</sup> | EffectorP<br>3.0 (Y/N) | Nucleotide<br>sequence<br>polymorphism<br>between two<br>parents (Y/N) | Position on 0-1<br>reference genome | Mature<br>length<br>(amino<br>acids) |
| --- | --- | --- | --- | --- | --- |
| PTT_10000004 | Y | Y | Y | Chr1: 62,330-62,746 | 122 |
| PTT_10008974 | Y | Y | Y | Chr8: 1870138-1870635 | 147 |
| PTT_10008993 | Y | N | N | Chr8: 1949091-1950411 | 423 |
| PTT_10009006 | Y | N | Y | Chr8: 1979970-1982483 | 837 |

<sup>a</sup>Y = Yes, N = No

## Notes

### Competing Interest Statement

The authors have declared no competing interest.

